# Parasitism of trees by marmosets (Primates: Callitrichidae) enhance tree turnover

**DOI:** 10.1101/2020.02.29.971325

**Authors:** João Pedro Souza-Alves, Guilherme V. Barbosa, Renato Richard Hilário

**Author notes:** **Correspondence:** João Pedro Souza-Alves. Postgraduate Program in Animal Biology, Department of Zoology, Federal University of Pernambuco, 1235, Postal code 50670-901, Recife, Brazil.

## Abstract

We tested if gouging by *Callithrix jacchus* affects tree survival. The proportion of dead gouged trees in forest fragments was higher than the proportion of dead non-gouged trees, with larger effect on smaller trees. The number of holes did not affect tree survival. Parasitism of trees by marmosets may enhance forest turnover.

## 1. INTRODUCTION

In natural environments, each species interacts with many other species through competition, predation, mutualism, parasitism, among others. These ecological relationships are important for biological community structuring and for the maintenance of biodiversity (Howe & Westley, 1988). Marmosets (genera *Callithrix, Cebuella* and *Mico* spp.) are small-bodied primates that are widely distributed throughout South America (Digby, Ferrari & Saltzman, 2011) These primates are relevant for assuming different ecological roles, as seed dispersers (Figueiredo & Longatti, 1997; Silva, Verona, Conde & Pires, 2018), predators of small animals (Silva, Alvarenga & Boere, 2008; Hilário & Ferrari, 2010; Amora et al., 2014), and prey of larger animals Bezerra, Barnett, Souto & Jones, 2009; Albuquerque et al., 2014).

Marmosets may feed on fruits, invertebrates, small vertebrates, flowers, nectar, fungi and leaves however, plant exudates are usually a dietary staple (Digby et al., 2010; Hilário & Ferrari, 2010; Teixeira et al., 2016). Indeed, marmosets are morphologically adapted for obtaining and digesting plant exudates (Ferrari, Lopes & Krause, 1993; Eng, Ward, Vinyard & Taylor, 2009; Vinyard et al., 2009; Forsythe & Ford, 2011). They are capable of applying greater forces in their bites (Eng et al., 2009), present elongated lower incisor teeth, which are of the same size as their canines (Coimbra-Filho & Mittermeier, 1976; Coimbra-Filho, 1980), and their teeth enamel is thicker at the labial side and thinner at the lingual side (Rosenberger, 1978; Nagomi & Natori, 1986) compared to other callitrichids, creating a sharp structure that allows the marmosets to gouge trees in order to promote exudate flow. Lacher, Fonseca, Alves & Magalhães-Castro (1984) classified this behaviour as “parasitism of trees” and compared marmosets to “large aphids”. These authors, however, did not evaluate the possible negative effects of gouging behaviour on the trees. The concept of parasitism involves the consumption of an organism by another and frequently includes some harm to the host organism (Esch & Fernandez, 2013). Therefore, parasitism causes a reduction of growth, survival and/or reproduction of the host organism (Agnew, Koella & Michalakis, 2000). However, it remains to be demonstrated that marmosets cause harm to the trees through their gouging behaviour.

Here, we aim to test if gouging by common marmosets (*Callithrix jacchus*) affects the survival of trees in forest fragments in North-eastern Brazil. We hypothesized that the presence of holes gouged by marmosets increases the mortality of trees and that the greater the number of holes, the greater the probability of tree death. We also hypothesized that larger trees will be more parasitism-tolerant, presenting higher survival rates when gouged than smaller trees.

## 2. METHODS

### 2.1 Study sites

The study was conducted in four Atlantic Forest fragments located in North-eastern Brazil. The vegetation in the region is classified as lowland tropical rain forest, one of the Brazilian Atlantic forest phytophysiognomies, that occurs at 50-100 m altitude (Veloso, Rangel-Filho & Lima, 1991). The first study site was the Dois Irmãos State Park (DISP), a 384 ha protected area located in an urban area in the municipality of Recife, Pernambuco. The forest canopy is *ca.* 20 m tall, and emergent individuals of *Pera glabrata, Aspidosperma discolor*, and *Tapirira guianensis* can reach up to 30 m (Guedes, 1998). The tree density of the area is 215 trees/ha. The second site, Mata do Curado Wildlife Refuge (MCWR), is also a protected area located in the municipality of Recife and is composed of 106 ha urban Atlantic Forest. Even though it is a protected area, it has suffered from selective logging and deforestation. The area has a tree density of 780 trees/ha, a mean basal area of 24.7 m^2^/ha, an average canopy height of 11.2 m, and a high incidence of plant species from the Melastomataceae and Monimiaceae families (Lins-e-Silva & Rodal, 2008). The third site, the Recife Botanical Garden (RBG), is separated from the MCWR by a highway (BR 232) and is composed of 11 ha of Atlantic Forest. In this site, the basal area of trees is 18.5 m^2^/ha (Sousa Júnior, 2006), and tree density is 203 trees/ha. The most common tree species are *Helicostylis tomentosa, Parkia pendula, Dialium guianense*, and *Schefflera morototoni* (Sousa Júnior, 2006). The fourth study site was the Usina São José (USJ), a 340 ha fragment of relatively undisturbed forest (Alves-Araújo et al., 2008). This site is located approximately 40 km from the municipality of Recife. In this site, the tree density is 211 trees/ha, and the richest botanical families are Leguminosae, Myrtaceae, Melastomataceae, Malvaceae and Sapotaceae (Melo et al., 2011). All the four sites host populations of common marmosets.

### 2.2 Sampling procedure and data analysis

In order to verify the effects of marmoset gouging behaviour on tree mortality, we randomly implemented four transects (250 m × 5 m; 0.5-ha) in each forest fragment. Whenever possible, the transects were placed at least 500 m from one another. Within the transects, we recorded whether each standing tree was alive or dead and whether these trees had holes gouged by marmosets in their bark. We counted the holes below a height of 5 m on the bark of the gouged trees, given that we were not able to assess the number of holes reliably above this height. We assessed the diameter at breast height (DBH) of all gouged trees (alive or dead) with a diametric tape.

We used Fisher’s exact tests to assess whether the proportion of live and dead trees differed between gouged and non-gouged trees. We tested whether the number of holes per tree differed between study sites through an Analysis of Variance (ANOVA). In addition, we performed a binomial generalized mixed model (GLMM) to assess whether tree DBH or the number of holes per tree had an effect on the survival of the gouged trees, considering site identity as a random factor. All analyses were performed using the R software 3.6.1 (R Core Team, 2019) with an alpha value of 0.05. Mixed models were carried in the package MASS (Venables & Ripley, 2002).

## 3. RESULTS

Overall, we evaluated 1685 trees, from which 58 (3.4%) presented holes gouged by marmosets. In general, the mean DBH of gouged trees was 35.5 ± *SD* 9.1 cm (range: 13.7 - 59.2 cm). Among the gouged trees, the average DBH of the living trees was 36.0 ± *SD* 8.8 (range: 13.7 - 59.2 cm), and the DBH of the dead trees was 35.0 ± *SD* 9.0 (range: 18.5 - 54.8 cm). At the RBG, we did not record any gouged trees, which prevented the evaluation of the effects of gouging on the survival of the trees at this site. The proportion of gouged trees that were dead was significantly higher than the proportion of non-gouged trees that were dead at two sites (DISP and USJ - Table 1). Considering all the sites together, the proportion of gouged trees that were dead was also significantly higher (Table 1).

**TABLE 1.** Number of live and dead trees in the four study sites and whether the trees presented holes gouged by marmosets. The proportion of trees according to their gouged status is presented within parentheses. We also present the Fisher’s exact tests p-values assessing the effect of the gouging in the survival of the trees.

| Site | Gouging<br>Status | Live | Dead | *P* |
| --- | --- | --- | --- | --- |
| Recife Botanical Garden | gouged | 0 | 0 | - |
|  | non-gouged | 389 (0.96) | 17 (0.04) |  |
| Mata do Curado Wildlife<br>Refuge | gouged | 8 (0.80) | 2 (0.20) | 0.0688 |
|  | non-gouged | 398 (0.96) | 17 (0.04) |  |
| Dois Irmãos State Park | gouged | 11 (0.69) | 5 (0.31) | 0.0380 |
|  | non-gouged | 366 (0.88) | 49 (0.12) |  |
| Usina São José | gouged | 27 (0.84) | 5 (0.16) | 0.0043 |
|  | non-gouged | 380 (0.97) | 11 (0.03) |  |
| Total | gouged | 46 (0.79) | 12 (0.21) | 0.0002 |
|  | non-gouged | 1533 (0.94) | 94 (0.06) |  |

Among the non-gouged trees, the proportion of dead trees was similar between sites (0.03 to 0.04), differing only in DISP (0.12 - Table 1). Conversely, among the gouged trees, the proportion of dead trees was more variable between sites (0.16 to 0.31). Thus, gouging by marmosets increases the probability of tree death by 2.6 to 5 times and seems to depend on site-specific factors. The number of holes per tree also varied between sites (F_2,55_=4.725; p = 0.0128), being greater in MCWR (46.2 ± *SD* 31.0) than at the other sites (USJ: 21.6 ± *SD* 31.1; DISP: 18.7 ± *SD* 19.1). Considering only the gouged trees, the probability of tree death decreased with tree size (DBH) but was not affected by the number of holes (Table 2). While less than 60% of the smaller trees (DBH<20cm) survived when gouged, this proportion approached 100% of the larger trees (DBH>50cm - Figure 1).

**TABLE 2.** Results of the Binomial Generalized Mixed Model (GLMM) relating whether the tree was alive or dead with its diameter at breast height (DBH) and the number of holes gouged by marmosets. The study sites were considered as random factors.

|  | *Intercept* | *Residual* |
| --- | --- | --- |
| Standard deviation | 0.7197 | 0.9383 |

Fixed effects:
|  | *Coefficient* | *Standard Error* | *P* |
| --- | --- | --- | --- |
| Intercept | -2.380 | 1.466 | 0.111 |
| DBH | 0.092 | 0.042 | 0.033 |
| Number of holes | 0.025 | 0.025 | 0.308 |

**Figure 1.**
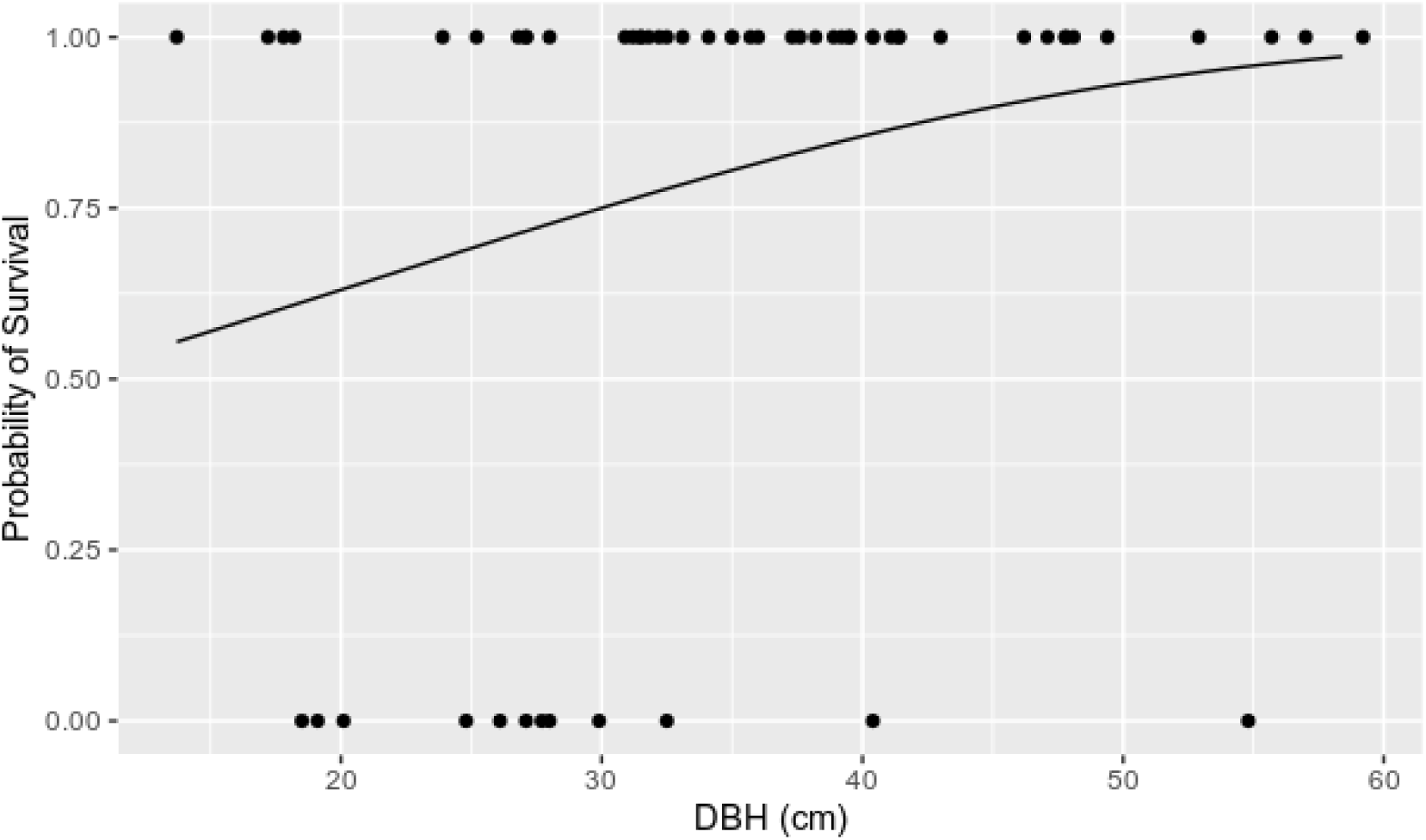
Probability of gouged tree survival according to their DBH (diameter at breast height), as accessed by a binomial model. The dots represent the live and dead trees sampled in the transects.

## 4. DISCUSSION

We showed that there is a larger proportion of dead trees among gouged trees, evidencing that gouging by marmosets increases the probability of death of the trees. This can be driven either by mechanical damage to the hydraulic system of the tree, interfering with the water supply (physiological stress), or by increasing the exposure of the tree to potential pathogens (Kautz, Meddens, Hall & Arneth, 2017; Tai et al., 2019). Furthermore, this effect is larger on smaller trees. It is likely that physical and mechanical factors, such as thinner barks, favour a greater relative damage on vessels (xylem and phloem) of smaller trees (Francisco et al., 2014), contributing to the increase in mortality. Finally, we confirmed the hypothesis proposed by Lacher et al. (1984) that marmosets may be classified as parasites of trees since they cause harm to the trees through their gouging behaviour.

Parasitism of trees represents an additional ecological role to marmosets and may be relevant to the forest dynamics since it should accelerate the turnover of the forest by increasing tree mortality (Stephenson & Mantgem, 2005). Also, since only some species are exploited for their exudates (Amora, Beltrão-Mendes & Ferrari, 2013; Thompson et al., 2013), parasitism may also influence interspecific competition among trees. This process may benefit pioneer trees, which are less-shade tolerant, fast-growing and less carbon-dense (Phillips, 1996), shaping forest composition and structure. Although the presence of holes gouged by marmosets increased tree mortality, the number of holes in the trees did not have such an effect. Marmosets usually concentrate their gouging on only a few individuals of some species, ignoring other tree individuals of the same species (Stevenson & Rylands, 1988; Thompson et al., 2013). Therefore, marmosets minimize the number of host individuals, reducing tree mortality. This behaviour may have been shaped by natural selection in order to avoid significantly reducing the availability of a key resource for the marmosets (i.e. plant exudates). Moreover, larger trees suffered less mortality from marmoset parasitism. Indeed, marmosets concentrate their gouging on larger trees (Lacher et al., 1984; Thompson et al., 2013), and this must be another strategy to reduce plant mortality as a result of tree parasitism.

We confirmed an additional ecological role for marmosets, as tree parasites, which may have effects on forest turnover. Over long periods, this process may substantially affect forest composition and structure, with possible effects on ecosystem services, such as carbon storage. By increasing tree mortality, the gouging behaviour of marmosets potentially reduce the availability of exudate sources for the marmosets. However, marmosets direct their gouging behaviour to a few large individuals, which should minimize their impacts on tree mortality. Although we studied only *C. jacchus*, our findings must also be true for other marmosets (genera *Callithrix, Cebuella* and *Mico*), and primates (i.e. *Phaner furcifer* and *Euoticus elegantulus*) that also gouge trees to some extent (Vinyard, Wall, Williams & Hylander, 2003; Forsythe & Ford, 2011).

## ACKNOWLEDGMENTS

We are grateful to all the staff of Dois Irmãos State Park, Usina São José, Mata do Curado Wildlife Refuge and Recife Botanical Garden. We also are grateful to Anielise Campêlo, Liany Oliveira-Silva, Ingrid Lima and Ana Caroline Araújo during the fieldwork. JPS-A was supported by Coordenação de Aperfeiçoamento de Pessoal de Nível Superior – Brasil (CAPES) - PNPD (Process no. 88882.306330/2018-1) and RRH was supported by CAPES - Edital 21/2018 (PROCAD-Amazônia - Process no. 88881.314420/2019-01). The study was also supported by FACEPE (BCT-0025-2.05-17).

## AUTHOR CONTRIBUTIONS

RRH and JPS-A designed the experiment, GVB collected the data, RHH analysed the data, and JPS-A and RRH wrote the manuscript. All authors approved the publication of this manuscript and agreed to be held accountable for the work performed therein.

## DATA AVAILABILITY STATEMENT

Data available from the Dryad Digital Repository: http://doi:10.5061/dryad.fxpnvx0nq (Souza-Alves, Barbosa & Hilário, 2020).

## ETHICAL GUIDELINES

All research complied with Brazilian legal requirements. It also adhered to the ASAB/ABS Guidelines for the Use of Animals in Research and American Society of Primatologists Principles for the Ethical Treatment of Non-Human Primates.

